# The Epstein-Barr virus episome maneuvers between nuclear chromatin compartments during reactivation

**DOI:** 10.1101/177345

**Authors:** Stephanie A. Moquin, Sean Thomas, Sean Whalen, Alix Warburton, Samantha G. Fernanadez, Alison A. McBride, Katherine S. Pollard, JJ L. Miranda

## Abstract

The human genome is structurally organized in three-dimensional space to facilitate functional partitioning of transcription. We learned that the latent episome of the human Epstein-Barr virus (EBV) preferentially associates with gene-poor chromosomes and avoids gene-rich chromosomes. Kaposi’s sarcoma-associated herpesvirus behaves similarly, but human papillomavirus does not. Contacts localize on the EBV side to OriP, the latent origin of replication. This genetic element, and the EBNA1 protein that binds there, are sufficient to reconstitute chromosome association preferences of the entire episome. Contacts localize on the human side to gene-poor regions of chromatin distant from transcription start sites. Upon reactivation from latency, however, the episome moves away from repressive heterochromatin and toward active euchromatin. Our work adds three-dimensional relocalization to the molecular events that occur during reactivation. Involvement of a myriad of interchromosomal associations also suggests a role for this type of long - range association in gene regulation.

**IMPORTANCE:** The human genome is structurally organized in three-dimensional space, and this structure functionally affects transcriptional activity. We set out to investigate whether a double stranded DNA virus, Epstein-Barr virus (EBV), uses similar mechanisms as the human genome to regulate transcription. We found that the EBV genome associates with repressive compartments of the nucleus during latency and active compartments during reactivation. This study is advances our knowledge of the EBV life cycle, adding three-dimensional re-localization as a novel component to the molecular events that occur during reactivation. Furthermore, the data adds to our understanding of nuclear compartments, showing that disperse interchromosomal interactions may be important for regulating transcription.

## INTRODUCTION

The chromatin of a human interphase nucleus is structurally and functionally organized at multiple scales. Chromosomes are subdivided into topologically associated domains (TADs), structural genomic units characterized by sharp boundaries that promote long-range interactions within but not between different domains (1). TADs on average measure ∼200 kb each (2) and fold into discrete globules in three-dimensional space (3). Each TAD may be classified into one of two large groups that correlate well with traditionally defined active euchromatin and inactive heterochromatin based on histone modifications, gene density, and polymerase occupancy (2, 4, 5). Furthermore, TADs preferentially associate with other TADs of the same type within the same chromosome, resulting in two main compartments in the nucleus, termed compartment A and compartment B, which again roughly correspond to euchromatin and heterochromatin (2, 4, 5). These two functionally distinct partitions of each chromosome are physically separated in the nucleus (6). Assembly mediated by long-range interactions also compacts individual chromosomes so that each occupies a discrete globular space known as a chromosome territory (7). These territories similarly partition into two main groups: gene-rich chromosomes found in the center of the nucleus and gene-poor chromosomes found at the periphery (8). The organization of DNA does not necessarily proceed in a hierarchal manner but nonetheless spans scales of many orders of magnitude. Clearly, genome structure and function are linked on many levels.

Interphase chromatin is highly dynamic in a manner coupled to changes in transcription. As the transcriptional state of a TAD changes, that unit rearranges intrachromosomal contacts to associate with similarly active or inactive domains (9). Reorganization is not, however, only restricted to changes in local interactions. Interchromosomal associations activate genes (10-12) and actively transcribed genes from different chromosomes colocalize (13, 14). In many cases, genes move out of a chromosome territory when activated (15, 16). Such long-range contacts make the spaces between chromosome territories a dynamic interface. Inhibition of transcription changes intermingling patterns of chromosome pairs, suggesting that gene activity may drive these associations (10). Taken together, these studies show that interchromosomal chromatin interactions are dynamic and that nuclear repositioning is often coupled with changes in transcription.

The human Epstein-Barr virus (EBV) is a double-stranded DNA herpesvirus that is maintained as an episome in the nucleus of a host cell. The viral genome is circular and chromatinized, resembling a small human chromosome in many molecular aspects. Like most herpesviruses, EBV establishes lifelong latency and occasionally undergoes spontaneous reactivation. The virus displays several different latent transcription programs in which different combinations of ∼10 or fewer transcripts are expressed. During reactivation, transcription drastically increases to ∼100 transcripts as the virus produces the proteins necessary for replication of the genome and packaging of new virions (17). Latent gene expression patterns are regulated by three-dimensional intrachromosomal interactions within the viral genome (18, 19), but how interchromosomal interactions between the virus and the human genome affect viral transcription is understudied for technical reasons.

Since the EBV genome has a similar structure to human chromatin and uses similar mechanisms to control transcription at the protein level, we wondered whether the virus also uses the 3D structure and functional organization of the nucleus to regulate gene expression and the genetic switch to a very transcriptionally active state. As a first step toward answering this question, we sought to understand the extent of engagement with this nuclear organization when double-stranded DNA viruses infect cells. Here we use in situ Hi-C (2) to measure interactions between the EBV genome and the human genome during latency and reactivation. We show that during latency, the EBV genome uses a small genetic element to interact with a network of repressive heterochromatin. Upon reactivation, the viral genome engages in different associations to leave this repressive environment and surround itself with active euchromatin.

## RESULTS

### Association of the EBV episome with the host genome depends on chromosome gene density

We measured spatial DNA-DNA colocalization with in situ Hi-C (2) to determine how the EBV episome interacts with human chromosomes and different nuclear compartments. To maximize signal for the transcriptionally quiescent form of the latent episome, we chose to examine Daudi, KemIII, RaeI, and Raji, Burkitt lymphoma cell lines that display very little spontaneous lytic reactivation capable of generating newly replicated liner genomes (20). Raji and Daudi cells contain ∼60 and ∼150 copies of the EBV episome, respectively (21). Our Hi-C data sets contain ∼17–40 million valid paired end contacts after quality control filtering, of which ∼4–9 million are interchromosomal and ∼10,000–230,000 between the EBV episome and human genome. We observe ratios of interchromosomal to chromosomal interactions indicative of high-quality experiments that detect proximity-dependent *in vivo* colocalization instead of non - specific *in vitro* artifacts (22).

First we examined interchromosomal interactions within the human genome at chromosome-level resolution by measuring observed interactions between different human chromosomes relative to random expectation. This metric normalizes for both random association and chromosome length. We calculated robust and high-confidence ratios by obtaining similar sequencing depth as the original Hi-C protocol that first measured chromosome-level interchromosomal interactions (5). The partitioning of gene-rich and gene-poor chromosomes into separate regions of nuclear space observed by Hi-C (5) further validate the original observations detected with fluorescence *in situ* hybridization (8). We ourselves detected small gene rich chromosomes such as 16, 17, 19, and 22 preferentially associating with each other, depicted by a cluster of red enriched nodes in the lower right of the heatmap (Fig 1A). We also detect the slightly weaker preferential association between large gene poor chromosomes such as 2–5, depicted by a second cluster of red enriched nodes in the upper left. The two groups of chromosomes tend to avoid one another as indicated by the cluster of blue depleted nodes in the upper right and lower left of the heatmap. We also note a few strong interactions that defy this pattern. For example, the artificially high interaction frequency between chromosomes 8 and 14 is due to the chromosomal translocation commonly found in Burkitt lymphoma (23) (Fig 1A, 1B, and 6A).

**FIG 1.**
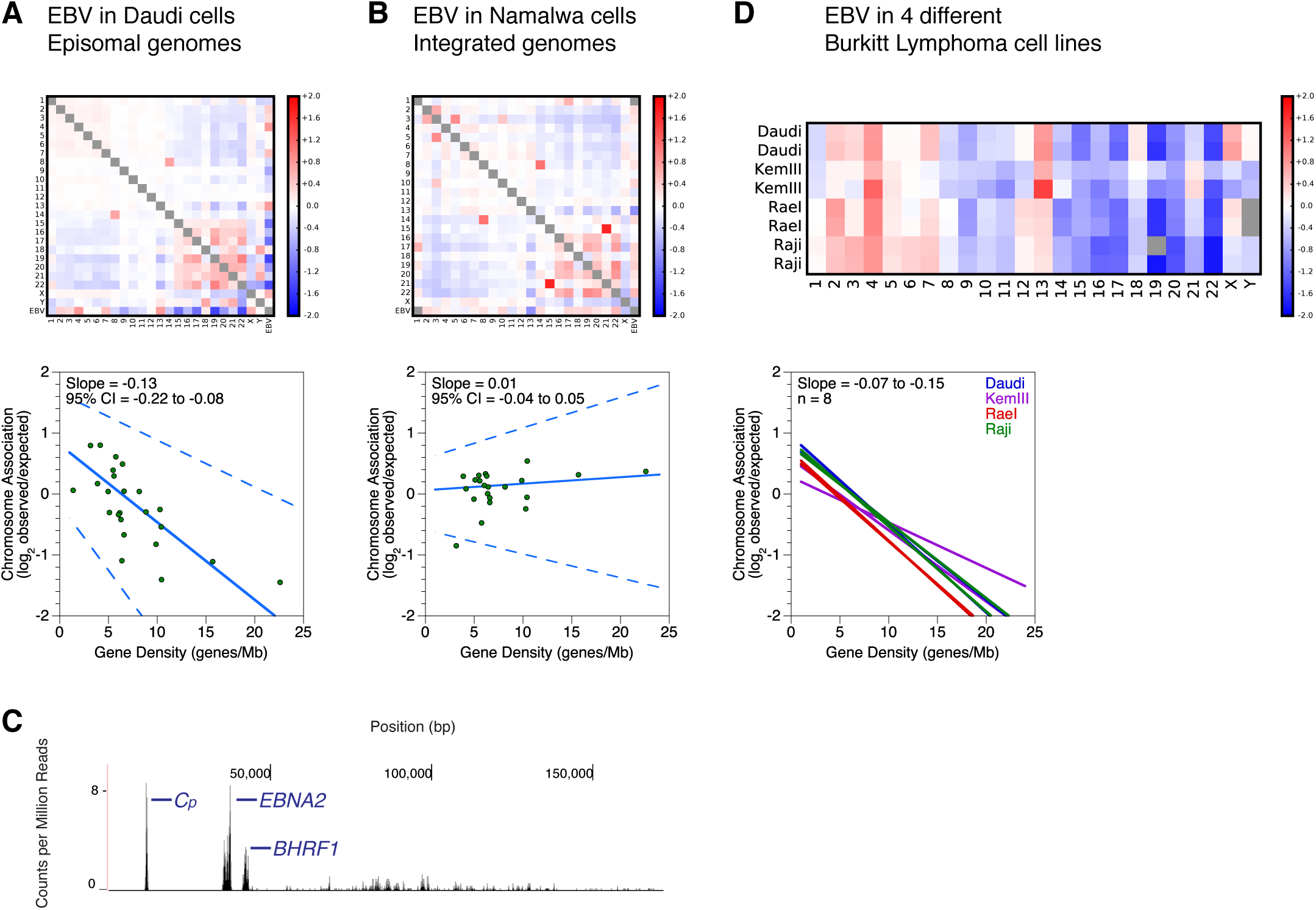
Episomal EBV genomes associate with the human genome in correlation with chromosomal gene density. (A, B) Interchromosomal contacts involving the EBV and human genomes in the Daudi and Namalwa cell lines measured by Hi-C. Heatmaps of chromosome association between chromosomes and between each human chromosome and the EBV genome. Observed counts are normalized against random expectation and shown on a log_2_ scale. Red indicates enrichment and blue indicates depletion. Below, scatterplots depict virus-human chromosome association plotted against gene density of each chromosome. Solid line indicates the Thiel-Sen fit and the dashed lines indicate the 95% confidence interval. Results are representative of two independent biological replicates. (C) Deep sequencing of EBV transcription in the Namalwa cell line. The X axis denotes nucleotide position and the Y axis denotes the number of counts per million mapped reads. RNA signals with unambiguously assignable annotations are marked. *BHRF1*, *Cp*, and *EBNA2* are latent transcripts labeled blue. Results are representative of two independent biological replicates. (D) Interchromosomal contacts between the EBV and human genomes in Burkitt lymphoma cell lines measured by Hi-C. Heatmaps of chromosome association between each human chromosome and the EBV genome in different Burkitt lymphoma cell lines. Shown are two replicates of four different cell lines: Daudi, KemII, RaeI, and Raji. Observed counts are normalized against random expectation and shown on a log_2_ scale. Red indicates enrichment and blue indicates depletion. Gray boxes represent either Y chromosomes not present in the female RaeI cell line or a score with absolute value greater than 2. Below, solid lines indicate the Thiel - Sen fit of virus-human chromosome association plotted against gene density of each chromosome. Each line represents one of two independent biological replicates for four different cell lines.

Next we examined interactions of the EBV episome with human chromosomes. Based on observed/expected chromosome association measurements, we discovered that the EBV episome avoids interaction with small gene rich chromosomes and preferentially interacts with large gene poor chromosomes (Fig 1A). This trend is conserved in the four cell lines we examined (Fig 1D). Although the exact order of the preferences varies slightly between cell lines, the strongest ratios are observed with the EBV episome avoiding chromosomes 16, 17, 19, 20, and 22 while interacting with 4 and 13. The observed trend does not correlate with size based on ratios calculated for chromosomes that defy the trend of gene density increasing as size decreases. The small but gene-poor chromosome 18 is not strongly avoided by the EBV episome; the large but gene-rich chromosome 1 is not strongly interacting with the EBV episome. EBV latency type, which varies between type I for RaeI and Daudi compared to type III for KemIII and Raji (20), also has no effect. To illustrate the suspected trend with statistical rigor, we plotted the observed/expected chromosome association preferences for EBV against gene density for each chromosome and calculated median slopes and 95% confidence intervals (CI) using Thiel-Sen linear regression (24). We found a negative slope in all Burkitt lymphoma cell lines (Fig 1D). The 95% CI completely falls in the negative range for all replicates (Fig 1A), demonstrating that the propensity of EBV to interact with a chromosome is strongly negatively correlated with the gene density of that chromosome.

### Preferential EBV chromosome associations require episomal genomes

To test whether viral chromosome preference is dependent on genome sequence or biophysical mobility, we performed in situ Hi-C on a cell line with integrated EBV. Namalwa cells contain an EBV genome with a sequence similar to others we examined, but it is integrated into chromosome 1. Gene expression predominantly consists of latent transcripts as measured by our RNA-seq experiments (Fig 1C), somewhat similar to the other Burkitt lymphoma lines studied. We found that integrated EBV does not show the same chromosome association preferences as episomal EBV and instead shows no correlation with gene density (Fig 1B), demonstrating that the viral genome must be episomal and less restricted to move around the nucleus in order to associate with chromosomes based on gene density. Furthermore, we note that the chromosome association preferences of integrated EBV in the Namalwa cell line are similar to chromosome 1, suggesting that the local chromatin environment at the integration site cannot be overcome by the viral sequence.

### Chromosome association preferences are conserved among some but not all episomal viruses

We also performed in situ Hi-C on three cell lines containing other latent double-stranded DNA viruses to determine whether chromosome association preferences are a conserved feature of episomal vectors. Two lines contained different strains of human papillomavirus (HPV), HPV16 and HPV31, which are unrelated to EBV, and one line contained the Kaposi’s sarcoma - associated herpesvirus (KSHV), which is closely related to EBV. We found that while KSHV does show a chromosome association preference similar to EBV, HPV16 and HPV31 do not (Fig 2). For KSHV, 95% CIs of calculated linear regression slopes fall within negative values, fitting a trend of lower chromosome association as a function of gene density. For HPV, regression slopes are close to 0 with the 95% CIs ambiguously spanning both negative and positive values. This demonstrates that while gene-density driven chromosome association preferences are not characteristic of all episomes, closely related gammaherpesviruses such as KSHV and EBV may use similar mechanisms to control their nuclear localization.

**FIG 2.**
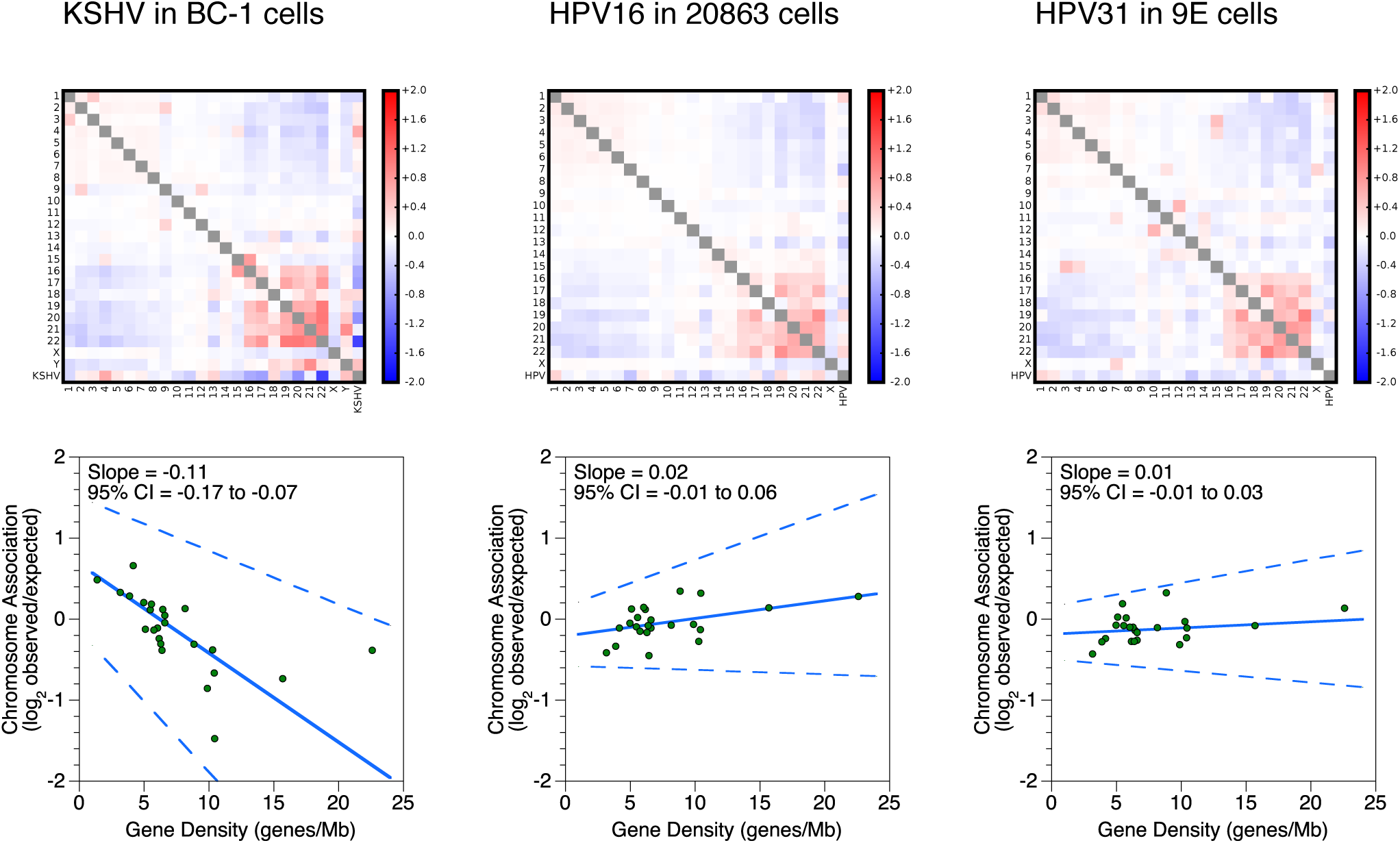
KSHV but not HPV genomes associate with the human genome in correlation with chromosomal gene density. Interchromosomal contacts involving the KSHV, HPV, and human genomes in the BC-1, 20863, and 9E cell lines measured by Hi-C. Heatmaps of chromosome association between chromosomes and between each human chromosome and the KSHV or HPV genome. Observed counts are normalized against random expectation and shown on a log_2_ scale. Red indicates enrichment and blue indicates depletion. Scatterplot shows the correlation between gene density of each chromosome and the virus-human chromosome association. Solid line indicates the Thiel-Sen fit and the dashed lines indicate the 95% confidence interval. Results are representative of two independent biological replicates.

### OriP and EBNA1 sufficiently reconstitute preferential EBV chromosome associations

To obtain a higher resolution understanding of which regions in the viral genome contact the human genome, we analyzed a publicly available Hi-C data set of EBV-infected B cells where sequencing depth far exceeds any other experiment to date. The Hi-C data set obtained with GM12878 cells (2) contains ∼5 billion pairwise contacts. We would have liked all of our experiments to have been performed with similar sequencing depth as the GM12878 data set, but this case represents an exceptional example in the field and is not tractably reproducible for financial reasons. We therefore complemented our own low-resolution experiments with the available high-resolution data. Our reanalysis that included the EBV genome sequence shows that viral chromosome preferences in this lymphoblastoid cell line are similar to that observed in Burkitt lymphoma lines (Fig 3A). We then measured contact frequencies between the EBV episome and human chromosomes and identified 117 significant interactions using the GOTHiC algorithm (25). Of these, 61 or 52.1% involved the 8-9 kb bin of the viral genome (Fig 3B), which falls into a viral cis-regulatory element called OriP, genetically defined as bp 7315–9312.

**FIG 3.**
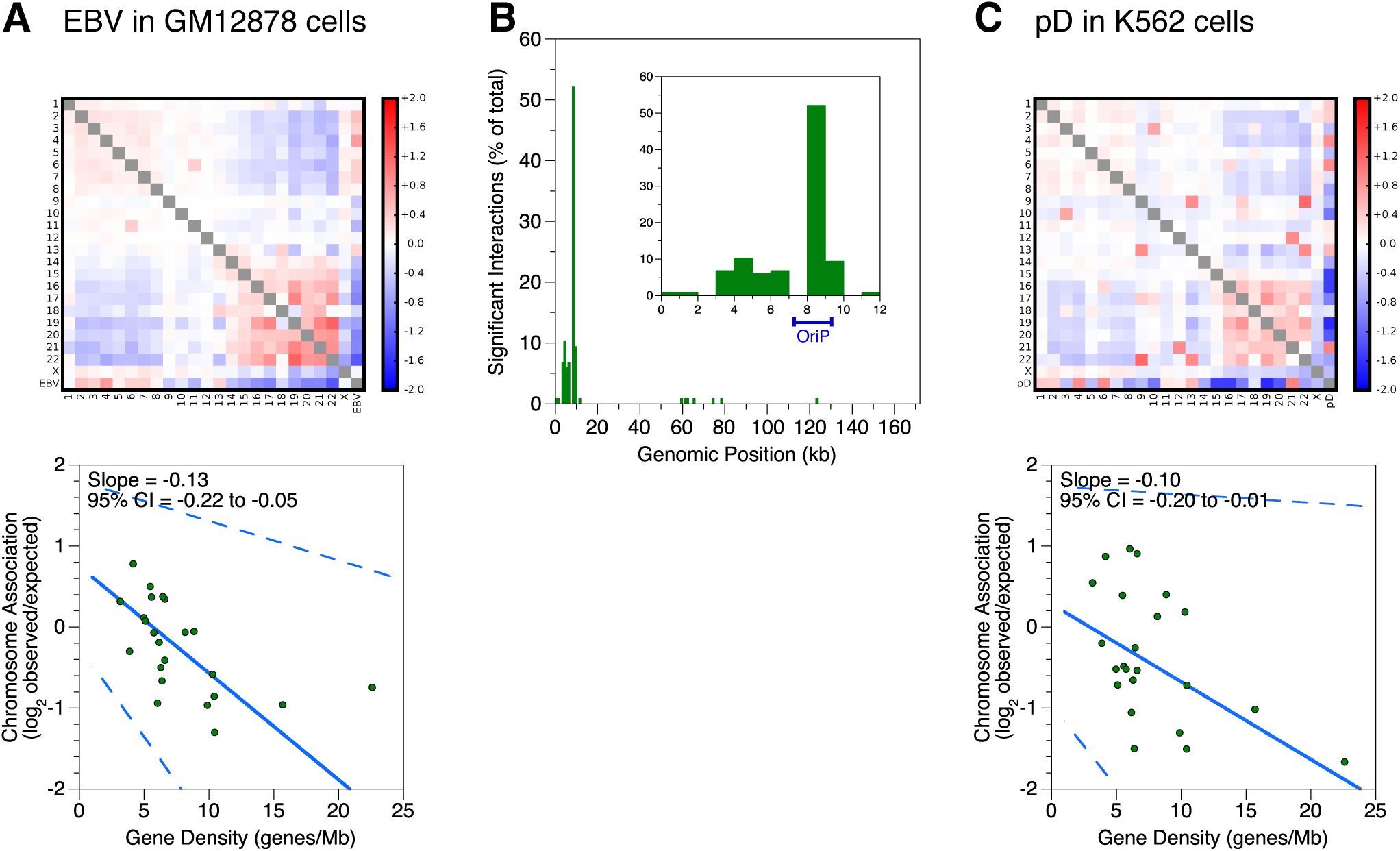
OriP is sufficient to reconstitute chromosome association preferences of full - length EBV. (A, C) Interchromosomal contacts involving EBV genomes, the pD plasmid, and human genomes in the reanalyzed GM12878 data set and K562 cell line measured by Hi-C. Heatmap of chromosome association between chromosomes and between each human chromosome and the EBV genome or pD plasmid. Observed counts are normalized against random expectation and shown on a log_2_ scale. Red indicates enrichment and blue indicates depletion. Scatterplot shows the correlation between gene density of each chromosome and the plasmid-human chromosome association. Solid line indicates the Thiel-Sen fit and the dashed lines indicate the 95% confidence interval. Results are representative of two independent biological replicates. (B) Localization within the EBV genome of significant interchromosomal contacts involving human chromosomes in the reanalyzed GM12878 data set. Histogram of unique significant interactions between human chromosomes and the EBV episome. The percentage of total interactions are plotted against position on the EBV genome in 1 kb bins. Inset depicts a zoom view of the first 10 kb of the viral genome.

To illuminate molecular properties of these intergenome loops, we applied the TargetFinder algorithm (26), which was previously used to determine features that predict long-range intrachromosomal interactions. We extended TargetFinder to model interchromosomal loops by requiring that one region from each interacting pair of segments is in the human genome while the other is in the EBV genome. Features used to predict interchromosomal interactions accordingly included genomic marks on the human or EBV segment. Perhaps surprisingly, histone modification ChIP-seq signals did not significantly contribute to accurate modeling of interchromosomal loops. Known chromosome organizers such as CTCF and cohesin also do not contribute significantly. The top predictive feature on the viral episome is occupancy by the viral protein EBNA1. Identification of this feature is consistent with the fact that EBNA1 binds OriP to perform two functions: replication of circular episomes during S phase as well as tethering of the viral genome to human chromosomes during metaphase (27). However, it is unknown whether EBNA1 mediates interactions between the viral episome and the human genome during interphase.

Considering that the majority of the significant interactions involved OriP, and that EBNA1 was a top predictor for significant human-viral contacts, we hypothesized that this assembly may be involved in interphase localization of the virus. To determine if OriP DNA and the EBNA1 protein are sufficient to reconstitute the chromosome association preferences of the entire virus, we performed in situ Hi-C on stably-transfected K562 cells containing pEBNA-DEST (pD), a plasmid that contains these two components. We found that pD showed similar preferences to the entire virus (Fig 3C), demonstrating that OriP and EBNA1 are sufficient to reconstitute preferential interactions with human chromosomes as seen with the full-length virus. Initial tests, however, imply that EBNA1 does not mediate chromosome preferences. We used shRNA to knockdown EBNA1 in RaeI cells, which contain full-length virus. We did not see a change in chromosome preferences upon ∼60% knockdown (Fig 4A). We also genetically deleted *EBNA1* from pD to generate pDΔEBNA and transiently transfected both vectors into K562 cells. Transient tranfections were necessary because *EBNA1* deletion precludes establishment of a stably maintained episome. The experiment is technically more variable and requires greater sequencing depth. Nonetheless, we still did not see a difference in chromosome preferences upon deletion (Fig 4B). This preliminary data suggests that a protein other than EBNA1 binds to OriP to mediate these preferences.

**Fig 4.**
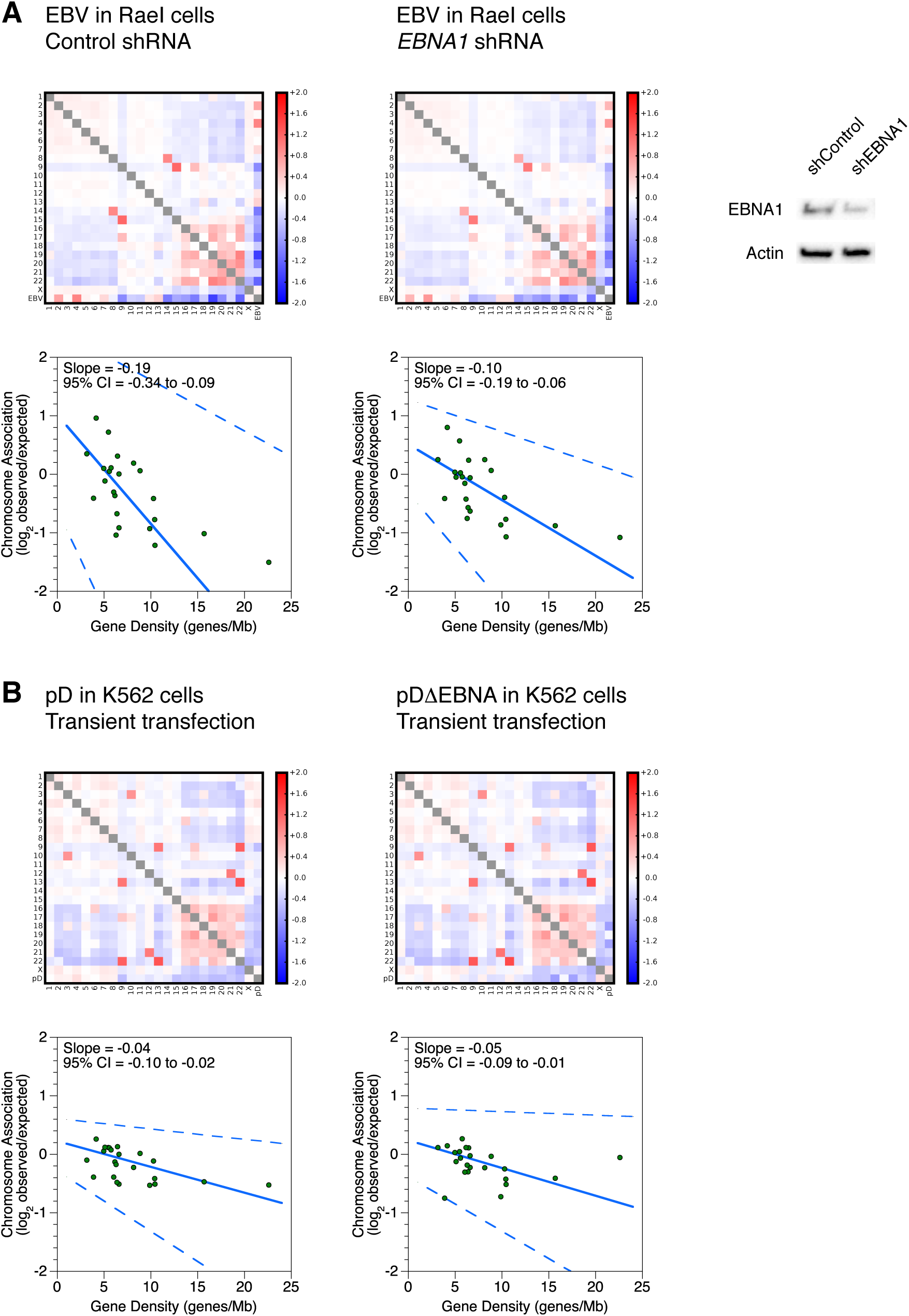
EBNA1 is not necessary to reconstitute chromosome association preferences of full-length EBV. (A, B) Interchromosomal contacts involving the pD plasmid, pDΔEBNA plasmid, EBV genomes, and human genomes in the RaeI and K562 cell lines measured by Hi-C. Heatmap of chromosome association between chromosomes and between each human chromosome and the EBV genome, pD plasmid, or pDΔEBNA plasmid. Observed counts are normalized against random expectation and shown on a log_2_ scale. Red indicates enrichment and blue indicates depletion. Scatterplot shows the correlation between gene density of each chromosome and the plasmid-human chromosome association. Solid line indicates the Thiel-Sen fit and the dashed lines indicate the 95% confidence interval. A gray box off the diagonal represents a score with absolute value greater than 2. (A) Lentivirus-mediated shRNA depletion of the EBV EBNA1 protein in the RaeI cell line. Western blots depict EBNA1 and β-actin expression levels in whole cell lysates after control or EBNA1 knockdown. (B) Deletion of the EBV EBNA1 gene in the pDΔEBNA plasmid in the K562 cell line. pD and pDΔEBNA are transiently transfected prior to measurement of interchromosomal contacts by Hi-C.

### EBV interacts with gene-poor human chromatin distant from transcription start sites

To characterize the human side of chromatin interactions with latent EBV, we studied the genetic landscape of enriched sites. We again used the significant contacts from the GM12878 data set filtered through GOTHiC. The 117 identified interactions localized to 91 unique 100 kb bins of human chromosomes. We used the Genomic Regions Enrichment of Annotations Tool (GREAT) (28) to measure gene density and distance to transcription start sites (TSSs). We compared the 91 significant regions to 100 sets of 91 randomly generated non-significant regions. Regions of the human genome that interact with the virus have ∼20% lower gene density, empirical p=0.18 (Fig 5A). Regions that interact with EBV are also farther from TSSs (Fig 5B). TSSs appear ∼20% less frequently within 500 kb, empirical p<0.01, and ∼250% more frequently beyond 500 kb, empirical p<0.01. We also used TargetFinder to ask which features were most predictive of viral-human interactions on the human side. We found that relative Hi-C coverage is the most predictive feature. In other words, regions of the human genome that interact with the viral episome have much higher Hi-C interaction frequencies in general compared to non-significant regions (Fig 5C). This result, however, does not imply some sort of non-specific technical artifact in our detection algorithm. We emphasize that peak-calling by the GOTHiC algorithm normalizes for sequencing depth. Although the highest coverage portions of the human genome colocalize with EBV, these regions still do so with a frequency higher than random expectation. Together, our analysis argues that EBV preferentially associates with interactive gene-poor regions of human chromatin far from TSSs.

**Fig 5.**
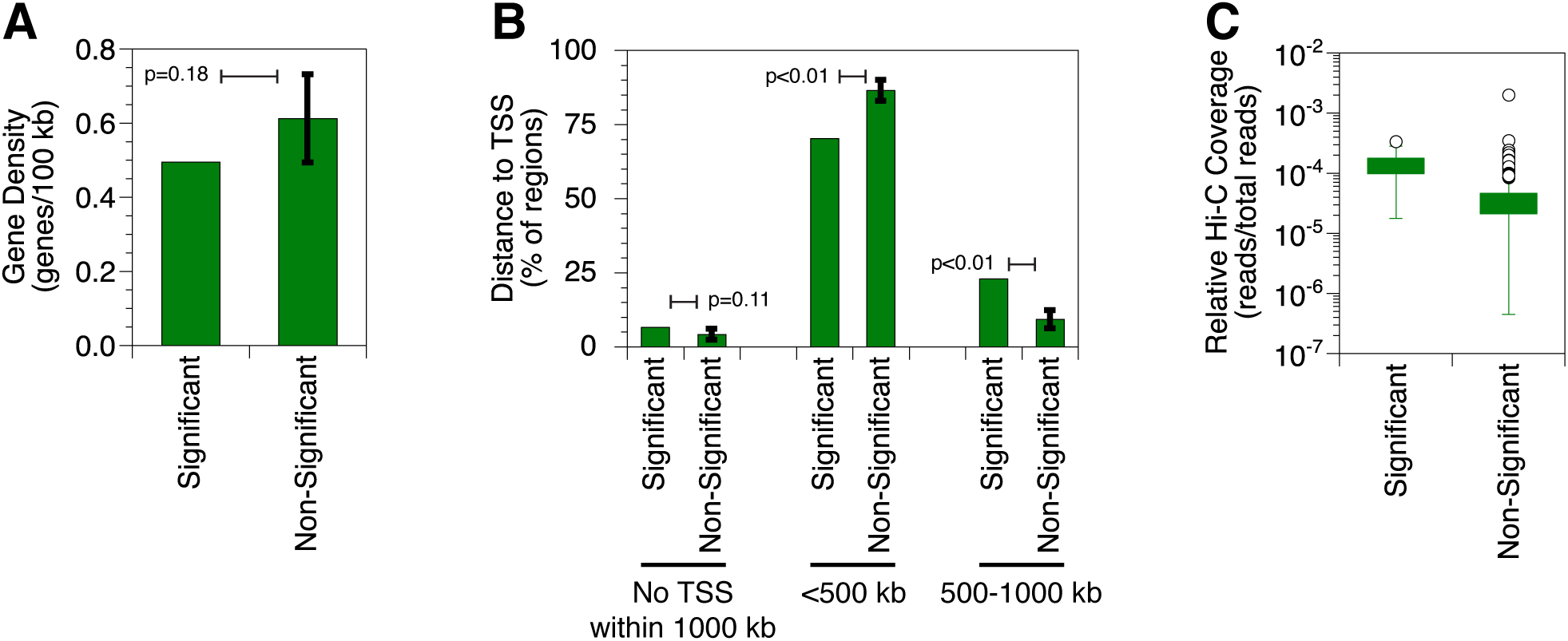
EBV episomes contact gene-poor human chromatin distant from transcription start sites. (A) Gene density of human genome regions that interact with the EBV episome in the reanalyzed GM12878 data set. The single set of significant interacting regions are compared to randomly chosen and equally large subsets of non-significant interacting regions. Error bar represents the standard deviation of 100 replicates resulting in an empirical p-value. (B) Distance to nearest TSS from human genome regions that interact with the EBV episome in the reanalyzed GM12878 data set. The single set of significant interacting regions are compared to randomly chosen and equally large subsets of non-significant interacting regions. Error bars represent the standard deviation of 100 replicates resulting in an empirical p-value. (C) Gene density of human genome regions that interact with the EBV episome in the reanalyzed GM12878 data set. Box and whisker plots for relative Hi-C coverage of human genome regions that interact significantly with the EBV genome compared to background. All significant interacting regions are compared to all non-significant interacting regions. Each box depicts 50% of the data. Whiskers extend 150% of the interquartile distance from the upper and lower quartiles with outliers shown as circles.

### The EBV episome switches associations from human heterochromatin to euchromatin during reactivation

A single snapshot of nuclear organization is often incomplete, so we measured whether interactions between the viral and human genome were remodeled based on changes in transcription. To do so, we compared in situ Hi-C results between cells containing the latent viral episome, which expresses ∼1-10 transcripts, and cells containing the lytic viral episome, which expresses ∼100 transcripts (17). We used the Akata-Zta cell line, which contains the EBV genome and an additional plasmid with a doxycycline-inducible promoter that produces the viral protein BZLF1, which induces lytic gene expression, as well as non-functional LNGFR, which facilitates purification of reactivated cells (29). We pre-treated Akata-Zta cells with acyclovir to block viral replication and ensure that we were examining interactions of episomes and not newly replicated linear genomes. DNA deep sequencing of acyclovir - and doxycycline-treated LNGFR - cells, which represent the latent population, estimate a viral genome copy number of ∼20. Acyclovir indeed functions as intended; total EBV DNA content increased by only 2.1±0.4-fold, much of which comes from abortive replication attempts near the lytic origins (30). Comparison of EBV episomes in latent and lytic cells revealed that the chromosome association preferences are lost during reactivation (Fig 6A and 6B). In lytic cells, linear regression slopes are close to 0 instead of strongly negative. This loss of preferential association based on gene density was reproduced in five independent reactivation experiments (Fig 6C).

**FIG 6.**
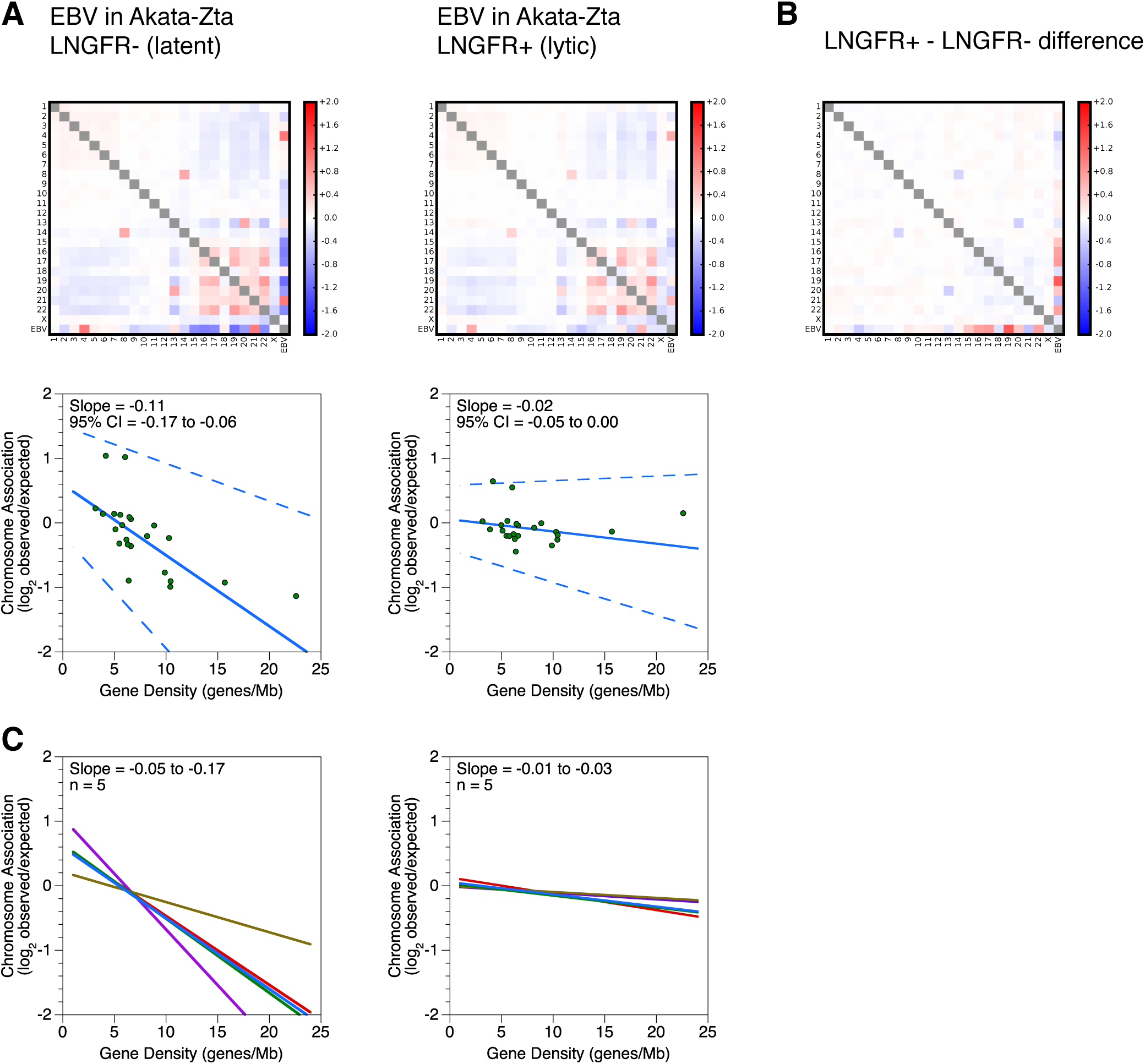
Chromosome association preferences of EBV episomes restructure during reactivation. (A) Interchromosomal contacts involving the EBV and human genomes in the Akata-ZTA cell line measured by Hi-C. LNGFR - and LNGFR+ cells contain latent and lytic episomes, respectively. Heatmaps of chromosome association between chromosomes and between each human chromosome and the EBV genome during latency and reactivation. Observed counts are normalized against random expectation and shown on a log_2_ scale. Red indicates enrichment and blue indicates depletion. Below, scatterplots depict virus-human chromosome association plotted against gene density of each chromosome. Solid line indicates the Thiel-Sen fit and the dashed lines indicate the 95% confidence interval. Results are representative of five independent and paired biological replicates. (B) Changes in interchromosomal contacts involving the EBV and human genomes in the Akata-ZTA cell line upon reactivation measured by Hi-C. Subtraction heatmap depicting differences between latent and lytic chromosome associations shown in (A). Chromosome association values during latency were subtracted from values during reactivation. Results are representative of five independent and paired biological replicates. (C) Interchromosomal contacts between the EBV and human genomes in the Akata-ZTA cell line measured by Hi-C. LNGFR - and LNGFR+ cells contain latent and lytic episomes, respectively. Solid lines indicate the Thiel-Sen fit of virus-human chromosome association plotted against gene density of each chromosome. Each line represents one of five independent biological replicates. Paired comparisons are matched by color.

Since gene-poor chromosomes contain more heterochromatin than gene-rich chromosomes, we hypothesized that the preferential chromosome association is due to EBV localizing with specific types of host chromatin. We subsequently surmised that this interaction would change upon reactivation of the virus. For bioinformatic analysis, we used lamin-associated domains (LADs) mapped by DNA adenine methyltransferase identification (31) as markers of heterochromatin. Different pieces of DNA in the same chromatin domain generally establish similar sets of long-range associations detectable by in situ Hi-C. We therefore classified bins of the human genome as either heterochromatin or euchromatin, calculated a characteristic interaction pattern for each type, and used logistic regression to determine which of these two classes the EBV genome more closely resembles. This algorithm differs from previous approaches (2, 5, 9) in that we explicitly consider interchromosomal interactions with the human genome instead of only intrachromosomal interactions. We therefore incorporate the role of the previously understudied yet prelavent associations between chromosomes and, in this case, genomes. Since heterochromatin is a transcriptionally repressive environment, we hypothesized that if viral transcription increases, association with repressive heterochromatin would decrease. We indeed found that when compared to viral genomes in latent cells, episomes in reactivated cells showed a strong decrease in interactions with LADs (Fig 7), suggesting that the episome is leaving the transcriptionally repressive environment of heterochromatin and moving toward the transcriptionally permissive environment of euchromatin during reactivation.

**FIG 7.**
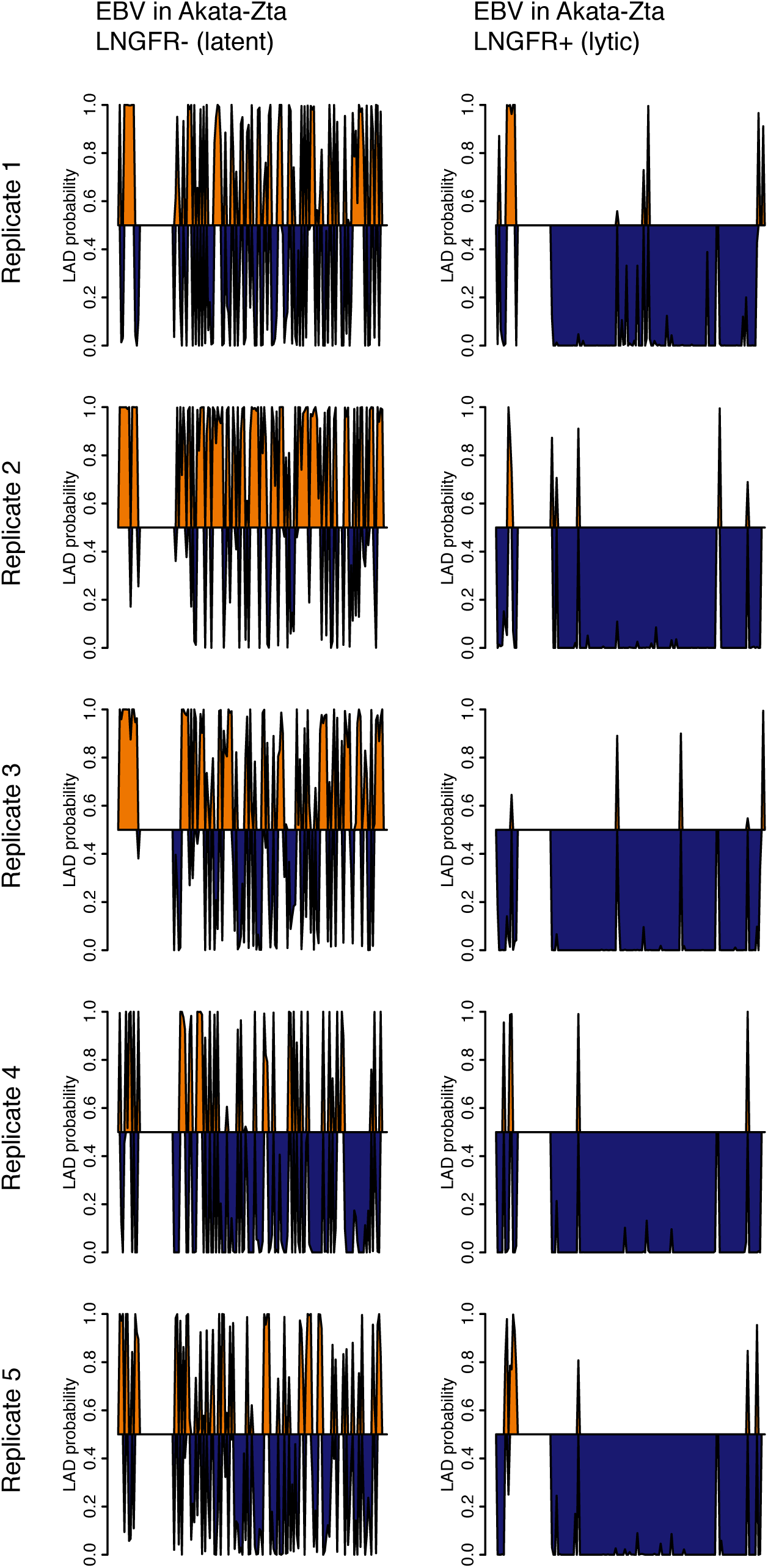
Predicted association of the EBV genome with LADs during latency and reactivation in the Akata-ZTA cell line. LNGFR - and LNGFR+ cells contain latent and lytic episomes, respectively. The probability of LAD association is plotted against position on the viral genome divided into 1 kb bins. Probabilities greater than 0.5 are shaded orange; probabilities less than 0.5 are shaded blue. Shown are five independent and paired biological replicates.

## DISCUSSION

Advances in high-throughput chromosome conformation capture technologies have now allowed us to quantitatively measure molecular interactions between host chromosomes and episomal pathogen genomes. While intrachromosomal interactions are well studied, interactions between chromosomes receive little scrutiny on the genome-wide level because previous methods were not sensitive enough to do thorough analysis. The original Hi-C assay performs proximity ligation with isolated protein-DNA complexes in dilute solution outside the context of a cell (5), but the majority of interchromosomal associations detected result from spurious events instead of proximity-dependent intra-complex ligation (22). Recent improvements to the original the Hi-C method structurally favor intra-complex ligation and yield lower rates of spurious events either through physical tethering to a surface, known as tethered Hi-C (4), or by performing the reaction within an intact nucleus, known as in situ Hi-C, (2, 32). In this study, we used the improved signal to noise ratio of in situ Hi-C to examine the interactions between host and pathogen genomes during viral infection. We also leveraged the copy number and high density transcriptional switching of the EBV episome to detect behavior not readily seen with autosomal loci. Our experiments reveal insight into the three-dimensional chromatin context of viral gene regulation as well general principles about the interplay of transcription and nuclear organization.

Very little was known about positioning of the viral genome within the nucleus or association with host chromosomes in three-dimensional space. Examining latent episomes, one previous study used three-dimensional fluorescence *in situ* hybridization after hypotonic chromosome condensation to detect possible colocalization between EBV episomes and human laminB1 (33) That observation is consistent with our own molecular data. The same work showed EBV predominantly colocalizing with activating histone modifications, and to a lesser extent repressive modifications. While we did not examine localization with histone modifications directly, our Hi-C data detects interactions with repressive heterochromatin during latency, an apparent contradiction with the microscopy results. Our molecular Hi-C data is of higher resolution and also examined localization with better preservation of nuclear organization by avoiding possible artifacts resulting from the hypotonic treatment used to visualize chromatin for microscopy. Another study used live-cell imaging to track nuclear positioning of lytic viral genomes during and after DNA replication but not during latency (34).

Our work specifically measures associations between EBV episomes with host chromosomes, comparing interactions during latency and reactivation.

We now know that even episomal viruses “integrate” into the network of human gene regulation. Here, we show that although EBV does not covalently integrate into the human genome, the virus noncovalently intermingles with the compartmentalized three-dimensional structure of the folded human genome. We determined that these interactions are non-random; the latent EBV episome preferentially interacts with gene-poor chromosomes and avoids gene-rich chromosomes. At higher resolution, the EBV episome associates with gene-poor regions of human chromatin distant from TSSs. The chromosome preferences can be reconstituted by OriP and EBNA1 alone. Surprisingly, initial results suggest that removal of EBNA1 does not change chromosome preferences, implying that another protein may be responsible for meditating these contacts.

The preferential association with human chromosomes during latency is not limited to EBV. We observe similar association patterns with KSHV. The strategy is not universally used by all episomal viruses, however, as we do not detect preferential interactions between HPV and human chromatin. The cause of this distinction, perhaps rooted in different selective pressures, remains to be elucidated.

We add another layer of understanding onto the sequence of molecular events coupled with viral reactivation. Much of what was previously known about the transition of EBV transcription from latent to lytic involves binding of viral and host proteins to the episomal genome (19, 35). Here we show that the virus changes nuclear environments during reactivation, switching from interactions with heterochromatin during latency to interactions with euchromatin compartment during reactivation. As a result, in both the latent and lytic transcription states, the viral genome is surrounded with human chromatin of similar transcriptional activity.

The movement of an episome upon reactivation argues that reactivation can drive passage between chromatin compartments with changes in only a diffuse network of interchromosomal associations without strong intrachromosomal contacts. We know that transcriptional changes correlate with transitions between compartments (9). This compartment switching, however, only identified changes in the predominantly detectable intrachromosomal contacts because previous computational methods did not consider colocalization between chromosomes (2, 5, 9). Here, we show that the EBV episome changes compartments through changes in diffuse interchromosomal associations as transcription increases during reactivation. Previous examples of functional interchromosomal interactions involve single contacts between two regions (11, 12). In contrast, the EBV episome forms a myriad of associations with the heterochromatin compartment during latency and a different set of distributed associations with euchromatin after transcriptional activation. The sum of many interchromosomal interactions may therefore contribute to gene regulation.

Future studies should involve tethering the viral episome to specific compartments, though recreating the network of interchomosomal interactions detected by Hi-C will be difficult. These experiments could elucidate the functional role of nuclear localization in EBV gene regulation: forcing connections with euchromatin may induce reactivation, while anchoring connections with heterochromatin may promote latency. The role of EBV chromatin in directing or responding to nuclear localization also requires clarification. The challenge still remains to determine whether transcriptional changes drive diffuse interchromosomal colocalization or vice versa.

## MATERIALS AND METHODS

### Cell culture and plasmids

EBV-positive Daudi, KemIII, RaeI, and Raji cells were maintained under standard conditions (20). K562 (36), EBV - and KSHV - positive BC-1 (37), and EBV-positive Namalwa (38) cells were maintained in RPMI - 1640 with 25 mM HEPES and 2 g/L NaHCO_3_ supplemented with 10% (v/v) fetal bovine serum (Invitrogen) in 5% CO_2_ at 37 **°**C. EBV-positive Akata-Zta cells (29) were maintained in RPMI-1640 with 25 mM HEPES and 2 g/L NaHCO_3_ supplemented with 10% (v/v) Tet System Approved fetal bovine serum (Clontech).

The HPV16-positive 20863 (39) and HPV31-positive 9E (40) keratinocyte cell lines were grown in F-medium, 3:1 (v/v) F-12-DMEM, 5% fetal bovine serum, 400 ng/ml hydrocortisone, 5 μg/ml insulin, 8.4 ng/ml cholera toxin, 10 ng/ml epidermal growth factor, 24 μg/ml adenine, 100 U/ml penicillin, and 100 μg/ml streptomycin, in the presence of irradiated 3T3-J2 feeder cells, as described previously (41).

Irradiated feeder cells were removed by versene treatment before the cells were harvested.

K562 cells were stably transfected with the pEBNA-DEST (pD) plasmid (Thermo Fisher Scientific) by a Nucleofector II device (Lonza) as directed, using solution V and program T-016. One day after transfection, 200 μg/ml hygromycin B was added and pD-positive K562 cells were selected for until transfected cells outgrew control cells and subsequently maintained in 200 μg/ml hygromycin B.

The *EBNA1* promoter and majority of the coding region were deleted from pD to generate pDΔEBNA by cutting with BsgI, filling in overhangs, and ligating blunt ends. Successful construction was verified by restriction digest mapping. For transient transfections, pD or pDΔEBNA plasmids were delivered into K562 cells using Fugene (Promega) at a 4:1 Fugene:DNA ratio using 20 μg DNA per 1 million cells. At 3 days post-transfection, 5 million cells were collected for Hi-C.

### In situ Hi-C

In situ Hi-C was performed with 5 million cells per experiment as described (2) with slight modifications. After end-repair and washes, Dynabeads (Thermo Fisher Scientific) with bound DNA were resuspended in 10 mM Tris, 0.1 mM EDTA pH 8.0 and transferred to new tubes. Sequencing libraries were created from bound DNA using the Ovation Ultralow Library System V2 kit (NuGEN) with one modification. After adapter ligation, because DNA is still attached to the beads, water instead of SPRI beads was added to the reaction. DNA bound to the beads was purified with a magnet, washed, and the beads were resuspended in 10 mM Tris, 0.1 mM EDTA pH 8.0. After library amplification, SPRI beads were added as directed to purify the amplified DNA. Quantitation and size distribution of libraries were performed using the Bioanalyzer High Sensitivity DNA Kit (Agilent). 50 base paired - end reads were sequenced on a HiSeq (Illumina).

Once sequenced, paired reads were aligned to combined human/viral reference genomes with HiCUP version 0.5.0 using default parameters (42) to generate a set of interactions. We used the human hg19 sequence merged with the EBV NC_007605.1, KSHV NC_009333.1, HPV16 NC_001526.2, or HPV31 J04353.1 sequence. The HiCUP processing steps remove PCR duplicates as well as invalid read pairs including those that are self-ligated or map to identical or adjacent fragments. Only alignments with mapq scores equal to or greater than 30 were retained. Data sets contained ∼7–40 million valid paired end HiC contacts after quality control filtering, of which ∼2–20 million were interchromosomal and ∼400– 230,000 between human and viral or plasmid sequences.

### Analysis of interchromosomal interactions

Chromosome-resolution heatmaps of interactions were determined from the HiCUP-filtered interchromosomal Hi-C interactions in sam format. Expected interactions were calculated using the following equation for each chromosome pair, where *chrA* represents the number of single end interchromosomal reads containing chrA, *chrB* represents the number of single end interchromosomal reads containing chrB, *totalPairs* represents the total number of interchromosomal paired end reads, and *all* represents the total number of interchromosomal single end reads, which is equal to 2 x *totalPairs*:

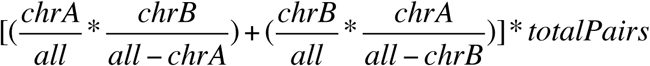

The chromosome association preference value for each combination was calculated by dividing the observed number of reads containing chrA and chrB by the expected value. Chromosome association preferences of viral genomes were plotted against gene density measured in genes per Mb (43). Data were fit to a line using the Thiel-Sen non-parametric linear regression median slope method (24) as implemented in the zyp R package.

### ChIP-seq

BZLF1 and EBNA1 chromatin immunoprecipitation deep sequencing (ChIP-seq) experiments were performed as previously described (30)using 3 μg of the anti-BZLF1 antibody BZ1 (Santa Cruz, sc-53904) and 3 μg of the anti-EBNA1 antibody 0211 (Santa Cruz, sc-57719).

### Analysis of viral-human contact regions

A large Hi-C data set from GM12878 cells (2) was reanalyzed to identify the strongest interactions between the EBV episome and human chromosomes. Chromosome-resolution heatmaps of interactions and chromosome association preference plots were generated from the HIC001 library. Further reanalysis included all fastq files from the primary and replicate sets, libraries HIC001–HIC0029. Data were independently processed using HiCUP version 0.5.8 and significant looping interactions were called using a Python implementation of the GOTHiC algorithm (25). Our specific parameters for performing the GOTHiC algorithm were as follows: To increase statistical power, reads were first mapped to fixed resolution bins, 100 kb on the human genome and 1 kb on the viral genome, using pairToBed as packaged in BEDTools version 2.26 (44). Counts from the two biological replicates were merged to increase the signal to noise ratio. Next, the probability of a spurious ligation was computed as a function of the relative coverage of each bin. Relative coverage was defined as the number of reads mapping to the bin divided by the total number of reads. Finally, the probability of observing a given number of reads by chance between two bins was computed using a binomial test, resulting in a p-value for each pair of bins. Multiple testing correction was performed using the Benjamini-Hochberg procedure, resulting in a q-value for each pair of bins. Bin pairs with a q-value of 0.1 or less, which corresponds to a 10% false discovery rate, were treated as statistically significant and identified as positive interacting samples.

Features that predict colocalization between the EBV episome and human chromosomes were identified using the TargetFinder algorithm (26). Average signals for all ENCODE GM12878 ChIP-seq data sets, as well as for the BZLF1 and EBNA1 ChIP-seq experiments performed by us, were computed for EBV and human bins. These features were used with interaction labels to train a gradient boosting classifier using the scikit-learn Python package (45) with the following parameters: n_estimators = 200, learning_rate = 0.01, and max_depth = 3. Stratified 10-fold cross-validation was performed with scikit-learn to obtain scores for precision, recall, and F1, the harmonic mean of precision and recall.

We characterized the genetic landscape of human chromosomal regions that colocalize with the viral episome with the Genomic Regions Enrichment of Annotations Tool (GREAT) (http://bejerano.stanford.edu/great/public/html/) (28). To measure gene density, the “basal plus extension” paramater was used, with a proximal extension of 50 kb upstream and 50 kb downstream, and a distal extension of 0 kb. Since GREAT chooses the midpoint of the 100 kb human bin as a reference, these settings allowed genes to overlap and permitted measurement of the total number of TSSs in each region. To determine distance to TSSs, the “single nearest gene” parameter was set to search within 1000 kb, allowing determination of the nearest TSS in either direction. Each of these two analyses were performed on the 91 significant bins, and on 100 sets of 91 random non-significant bins. An empirical p-value was measured to determine significance.

### shRNA-mediated EBNA1 knockdown

RaeI cells were transduced with shEBNA1 or control shRNA (46) by spinoculation. Lentivirus with 8 ug/ml polybrene was added to cells and spun at 800 xg for 30 minutes at room temperature. The supernatant was aspirated and cells resuspended in fresh media. After 48-72 hours post-transduction, RaeI cells were selected with 2 μg/ml puromycin. At 7 days post-transduction, cells were collected for Western blotting and 5 million cells were collected for in situ Hi-C.

Western Blots were performed using standard techniques. EBNA1 was detected using the anti-EBNA1 antibody 1EB12 (Santa Cruz, sc-81581) at 1:100-200 dilution and goat anti-rabbit-HRP (Abcam ab721) at 1:2,000-5,000 dilution. For normalization, actin was detected using an anti-β-actin antibody (Abcam, ab8227) at 1:10,000-20,000 dilution and rabbit anti-mouse-HRP (Abcam ab6728) at 1:20,000-30,000 dilution. Signals were detected using the SuperSignal West Pico Chemiluminescent Substrate (Thermo-Fisher), and a ChemiDoc MP Imaging System (BioRad). ImageLab (BioRad) version 5.2.1 was used to measure knockdown.

### Viral reactivation

Log-phase cultures of Akata-Zta cells were pretreated with 200 μM acyclovir (Sigma-Aldrich, A4669) for 1 hour before reactivation of the lytic cycle with 500 ng/mL doxycycline (Sigma-Aldrich). These cells contain a doxycycline-inducible plasmid with a bi-directional promoter that produces non - functional LNGFR along with the immediate early protein BZLF1, which starts the lytic cycle gene expression cascade and reactivates the virus (29). After 1 day, cells were magnetically sorted using LNGFR Microbeads and LS columns (Miltenyi Biotech).

EBV DNA quantitation was determined by deep sequencing total DNA. Genomic DNA was purified by silica-based membrane affinity with a DNeasy Blood & Tissue Kit (Qiagen). Libraries were constructed and the percentage of EBV reads from total measured (30). Viral genome copy number was estimated based on observed ratios between the number of reads mapped to the EBV and human genome sequences compared to the expected value, which is calculated as the ratio between the viral episome length and the summed length of all human chromosomes.

### LAD state predictions

The set of lamin interacting domains from Tig3 cells (31) was downloaded from the UCSC genome browser (47). Hi-C interactions identified by HiCUP were processed into a format readable by the HiTC package (48) using 1 Mb bins for the human genome and 1 kb bins for the viral genome. We performed further analyses on these interaction matrices in R (49). First a full interaction matrix for all autosomes was constructed. Next, for each bin, the mean of interaction counts with LAD bins was calculated as was the mean of interaction counts with non-LAD bins. This created two vectors of interaction means whose lengths were the number of bins in the autosomal genome. These vectors contained the primary source of information linking LAD state and interaction counts.

Correlations were then performed across all autosomal bins with the LAD mean vector and the non-LAD mean vector. A LAD bin will have a high correlation in interactions with the LAD mean counts and a low correlation in interactions with the non-LAD mean counts. A logistic regression was therefore performed on the LAD and non-LAD correlation values to estimate the probability with which each genome region would interact with lamin. Next, we calculated the probability that each viral bin interacted with the lamin by applying the logistic regression model of the autosomal LAD correlations to the viral interaction data. The R function and associated files used to perform this calculation are included with this manuscript (Supplementary File).

### Accession codes

Deep sequencing data was deposited in the Gene Expression Omnibus database under accession code GSE98123.

## SUPPLEMENTAL MATERIAL

### SUPPLEMENTARY FILE

ZIP file, 0.1 MB. R script for lamin state predictions and associated input bins files.

## ACKNOWLEDGMENTS

We are grateful to the Gladstone Genomics Core and the UCSF Center for Advanced Technology for use of shared equipment. We thank Paul M. Lieberman for the EBNA1 shRNA constructs. This project was supported the NIAID (R21AI108597 to J.L.M.), the Leukemia and Lymphoma Society (New Idea Award to J.L.M.), the NIMH (R01MH109907 to K.S.P.), the Bench to Bassinet Program of the NHLBI (UM1HL098179 to K.S.P.), the San Simeon Fund (K.S.P.), and the Gladstone Institutes (J.L.M. and K.S.P). This project was funded by the UCSF Program for Breakthrough Biomedical Research (New Frontier Research to J.L.M.), funded in part by the Sandler Foundation. This research was supported by the Creative and Novel Ideas in HIV Research Program (CNIHR) (J.L.M.) through a supplement to the University of California at San Francisco (UCSF) Center For AIDS Research funding (P30 AI027763). CNIHR funding was made possible by collaborative efforts of the Office of AIDS Research, the National Institute of Allergy and Infectious Diseases, and the International AIDS Society. A.W. and A.A.M. were funded by the Intramural Research Program of the NIAID, NIH. S.A.M. was supported by a US National Science Foundation Predoctoral Fellowship and the Microbial Pathogenesis and Host Defense training grant from the NIAID (T32AI060537).

## AUTHOR CONTRIBUTIONS

S.A.M., S.T., S.W., A.A.M., K.S.P., and J.L.M. designed research; S.A.M., S.T., S.W., A.W., S.G.F., A.A.M., and J.L.M. performed research; S.A.M., S.T., S.W., A.A.M., K.S.P., and J.L.M. analyzed data; S.A.M. and J.L.M. wrote the paper.

